# Greater alteration of gut microbiota occurs in childhood obesity than adulthood obesity

**DOI:** 10.1101/2022.05.22.492992

**Authors:** Zhongjia Yu, Xiang-Fang Yu, Xiu Zhao, Pei-Gen Ren

**Affiliations:** Center for Energy Metabolism and Reproduction, Shenzhen Institute of Advanced Technology, Chinese Academy of Sciences, Shenzhen, China; Shenzhen College of Advanced Technology, University of Chinese Academy of Sciences, Shenzhen, China; Shenzhen Children’s Hospital, Shenzhen, China

**Keywords:** children, adults, gut microbiota, obesity

## Abstract

The children’s gut microbiota, associated with the development of obesity, is in maturation. The impact of obesity on the gut microbiota in childhood could have a more significant effect than on adulthood and eventually be lifelong lasting, but it has been rarely studied. Aimed to discover the difference in gut microbiota between obese children and adults, we collected and re-analyzed published data from National Center for Biotechnology Information (NCBI). We found that the gut microbiota had a significant difference between children with and without obesity, but this did not happen in adults. Based on the random forest model, childhood obesity is more challenging to predict using gut microbiota than adulthood obesity. Our results suggest that gut microbiota in childhood is more easily affected than in adulthood. Hence, early intervention for childhood obesity is essential to improve children’s health and lifelong gut microbiota-related health.

## 1 Introduction

The association between gut microbiota and obesity has been widely discussed in the past decades. However, the etiologic relationship still lacks solid evidence for in-depth analyses^1^. Relevant studies focusing on both adulthood and childhood obesity have revealed that the shifts of gut microbiota happen during the development of obesity and during the intervention for obesity^2–5^. In adults, the signature of gut microbiota reflects the characteristics of the population in subgroups, including diet preference, living style, and risk factors for metabolic disorders^6^. Therefore, the signature of gut microbiota has become one of the most promising factors in predicting metabolic disorders, including obesity. Unlike adults, the gut microbiota of children is maturing; thus, the factors and status of childhood obesity may affect the shaping of gut microbiota and even have a lifelong impact^7^. However, minimal studies have been focused on the difference in gut microbiota between adulthood and childhood obesity. Moreover, cohorts from different countries or areas have their unique gut microbiota signatures due to the genetic factors, diet preference, and environment. These variants add difficulties in patterning the shifts of gut microbiota in the development of obesity worldwide. A study reviewing worldwide data to illustrate the different gut microbiota signatures in different countries would add value to understanding the relationship between obesity and gut microbiota. In the present study, we analyzed worldwide raw 16S rRNA amplicon sequencing data of gut microbiota in both adulthood and childhood obesity studies from the National Center for Biotechnology Information (NCBI) database to reveal different traits between obese adults and children in the world.

## 2 Materials and methods

### 2.1 Data resource

All raw data was obtained in the search with keywords “obesity” OR “obese” AND “gut microbiota” OR “microbiome” OR “microbiota” in the BioProject of NCBI. The data released before 1st February 2022 were searched. Only anthropic studies containing at least four healthy individuals were further analyzed. Raw data of 16S amplicon sequencing for each Biosample was collected. Moreover, if the study is about intervention, only basal data of both control and obese individuals were included.

All information on each BioProject was summarized using both published sample sheets (supplementary tables) and relevant publications. The category of bodyweight was in line with the sheets. If it was not referred, the following rules were used for BMI-grouping. For children from Non-US and US, the category of bodyweight was checked using BMI-for-ages check forms of World Health Organization and Centers for Disease Control and Prevention, respectively. For adults, the overweight is BMI between 24.9-30 kg/cm^2^, and obesity is BMI over 30 kg/cm^2^. After BMI-grouping, the category of bodyweight in the present study was redefined as the obesity included “overweight”, “obese,” and “superobese” from the BMI category, and other levels were counted as healthy control.

### 2.2 Data process

Raw data was analyzed using the QIIME2 (version 2021.08) software pipeline^8^. Reads were demultiplexed with q2-demux. Then, the DADA2 plugin was implemented for the quality control process, and all phiX reads and chimeric sequences were filtered. Based on the demux summary, single-end sequences were truncated to a length with 160-200 bases (details in supplementary table). After denoising, the data using the DADA2 denoise-paired method, representative sequences of each sample were retained. ASVs were filtered out non-bacterial sequences against Silva 99% using method vsearch, respectively. Then, Naive Bayes classifiers pre-trained on Silva 138 99% OTUs full-length sequences were used correspondingly for taxonomy assignment. Afterwards, the number of 16S rRNA copies of each taxon were normalized using a plug-in commend.

### 2.3 Statistic analyses and visualization

The principal coordinate analysis (PCoA), PERMANOVA test, and visualization were performed using the ImageGP online tool^9^. Alpha diversity index was calculated using QIIME2 plug-in commends and visualization using GraphPad. Differences in the abundance of each taxon between healthy and obese individuals were analyzed using DESeq^10^. The random forest model was tested using the R package^11^.

## 3 Results

A total of ten cohorts from nine BioProjects were included, four for childhood obesity (PRJNA647465^12^, PRJNA317290, PRJNA433269^13^, PRJNA637782^14^) and five for adulthood obesity (PRJNA483803, PRJNA339677^15^, PRJNA631293^16^, PRJNA273824, PRJNA417691^17^). Among these projects, 601 samples of raw data met our categories and were further analyzed, and the detail of each BioProject is shown in supplementary tables.

Regardless of ASVs or species level, there was no significant difference in gut bacterial composition between obese and healthy individuals in adults studies, except for one study of Mexican adults and ASVs level for Brazilian study (Fig.1A, supplementary Fig 1&2). For childhood obesity, all projects presented a significant difference in gut microbiota between children with and without obesity (Fig.1A, supplementary Fig 1&2). The age distribution is also shown in Fig. 1A. Except for the study for Mexican adults, all the studies compared control and obesity in the same age range. Alpha diversity index Shannon and Chao1 showed a trend that adulthood gut microbiota was more diverse than childhood ones, but was not significant (Fig 1B).

**Fig 1.**
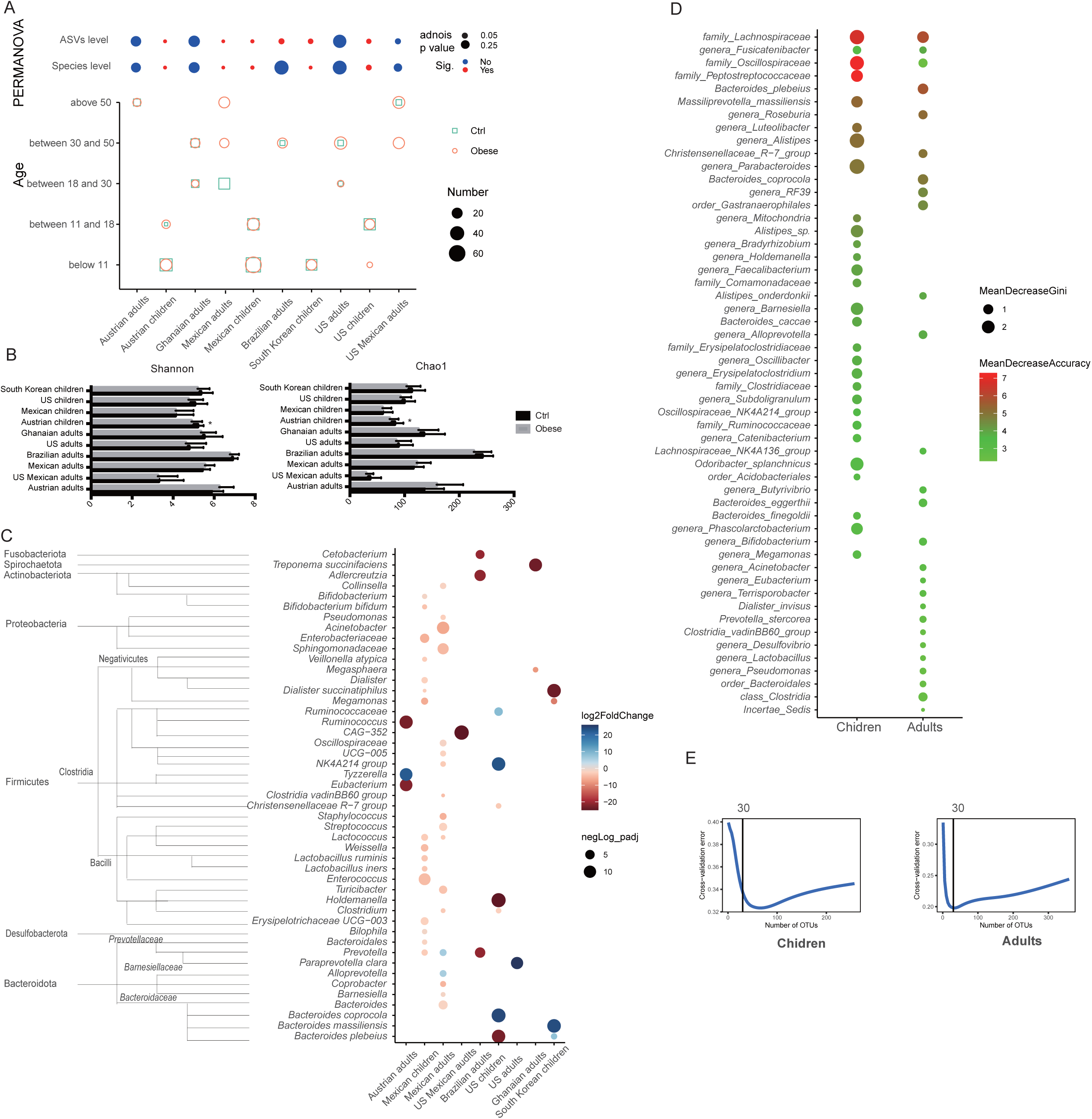
Comparison of gut microbiota between obese and control groups in different cohorts. A. Distance of gut microbiota between obese and control individuals analyzed using PERMANOVA and age distributions in different cohorts; B. Alpha diversity of gut microbiota in different cohort; C. Bacteria with significant difference in abundance between control and obese individuals in different cohorts (Blue means bacteria had higher abundance in control group than obese group, vice versa); D. Top 30 taxa used to predict obesity using the random forest model; E. Cross validation errors of both children cohorts and adults cohorts in random forest model.

Comparing the difference in abundance of bacteria between obese and non-obese groups, these differences were various depending on cohorts (Fig 1C). More bacteria belonging to Firmicutes and Bacteroidota had a significant difference in abundance, as the increase of Firmicutes and the decrease of Bacteroidota were generally observed in obese group. Except for the NK4A214 group of the family *Oscillospiraceae* and *Prevotella* presented different changes between different cohorts; other bacteria had consistent changes along with the obesity. For children in Austria, there were no bacteria observed a significant difference in abundance between children with and without obesity.

The random forest model was used for predicting obesity. As shown in Fig 1D, top30 taxa used to predict obesity in both children and adults had some overlaps. The family *Lachnospiraceae* is the top1 biomarker for obesity prediction with the highest accuracy. However, the general accuracy of using these taxa for the prediction is minimal, as there would be at least 20% for adults and 32% for children error generated in this prediction model. More details on each BioProject can be found in the supplementary tables.

## 4 Discussion

Childhood obesity is concomitant with the maturation of core gut microbiota and body development; hence it may have a lifelong influence on individuals. To prove if childhood obesity has a more significant impact on shaping gut microbiota than adulthood obesity, we collected relevant data worldwide in the public database as much as possible. We analyzed them with a uniform pipeline in the present study. In the same age ranges, children have more distinct gut microbiota between healthy and obese groups than adults. Meanwhile, higher error from the random forest model was generated when predicting childhood obesity than adulthood obesity, suggesting more difficulties in predicting childhood obesity correctly using gut microbiota. Moreover, based on the alpha diversity index, we could find a trend that gut microbiota in children was less diverse than in adults, indicating that the former has excellent potential to develop. Therefore, obesity and gut microbiota might be age-dependent. When obesity happens at a younger age, it has a more significant impact on gut microbiota shaping. It suggests that the childhood gut microbiota with obesity are quite different from healthy ones. Changes in the gut microbiota of children due to obesity may have a lifelong impact on the health of individuals, which should be more focused on public health.

Same as previous studies, we observed an increase in the ratio of Firmicutes to Bacteroidota in these cohorts^1,18^. More Firmicutes were found to increase significantly in obese groups and vice versa for Bacteroidota. However, specific bacteria with a significant difference in abundance between non-obese and obese individuals depended on cohorts. Commonly used probiotics, such as *Bifidobacterium* and *Lactobacillus*, only present significant differences in some cohorts, suggesting they might have minimal impact on other cohorts. Therefore, using one golden bacterium to prevent or treat obesity is challenging for all cohorts. The family *Lachnospiraceae*, a propionic acid producer that plays a crucial role in metabolic regulation^19^, showed its potential as a promising biomarker and metabolic regulator in the present study. In predication for childhood and adulthood obesity, the relative abundance of family *Lachnospiraceae* is the best biomarker.

However, only several cohorts were included due to the limited number of released data in the database. We would like to appreciate and encourage more contributions of sequencing data for further discovery. Moreover, the amplicon sequencing data used in the present study limits precise taxonomic classification. Therefore, tools with higher accuracy, such as metagenomics and culturomics, should be considered.

In conclusion, childrenhood obesity has a profound impact on shaping gut microbiota. More focus and earlier intervention for childhood obesity are crucial for the child’s health and lifelong gut microbiota-related health.

## Acknowledgement

Authors appreciate the contribution of raw data from incited projects. This work was supported by Shenzhen Fundamental Research Program (No. KCXFZ20201221173400002)

## Author Contributions

ZY and PR designed the study; ZY conducted data collecting, analyses and wrote the manuscript; PR, XY and XZ revised the manuscript.

